# Ff-nano, Short Functionalized Nanorods Derived from Ff (f1, fd or M13) Filamentous Bacteriophage

**DOI:** 10.1101/015511

**Authors:** Sadia Sattar, Nicholas J. Bennett, Wesley X. Wen, Jenness M. Guthrie, Len F. Blackwell, James F. Conway, Jasna Rakonjac

## Abstract

F-specific filamentous phage of *Escherichia coli* (Ff: f1, M13 or fd) are long thin filaments (860 nm x 6 nm). They have been a major workhorse in display technologies and bionanotechnology; however, some applications are limited by the high length-to-diameter ratio of Ff. Furthermore, use of functionalized Ff outside of laboratory containment is in part hampered by the fact that they are genetically modified viruses. We have now developed a system for production and purification of very short functionalized Ff-phage-derived nanorods, named Ff-nano, that are only 50 nm in length. In contrast to standard Ff-derived vectors that replicate in *E. coli* and contain antibiotic-resistance genes, Ff-nano are protein-DNA complexes that cannot replicate on their own and do not contain any coding sequences. These nanorods show an increased resistance to heating at 70 °C in 1 % SDS in comparison to the full-length Ff phage of the same coat composition. We demonstrate that functionalized Ff-nano particles are suitable for application as detection particles in sensitive and quantitative “dipstick” lateral flow diagnostic assay for human plasma fibronectin.

## 1. Introduction

Filamentous bacteriophages are filament-like bacterial viruses (Day, 2011;Rakonjac et al., 2011). The F-pilus-specific filamentous phage *of Escherichia coli*, Ff (f1, M13 and fd), are resistant to heat, wide range of pH and ionic detergents, to which the tailed phage, such as λ, are sensitive (Branston et al., 2013). The Ff phage virions are 860 nm in length and 6 nm in diameter; they contain circular single-stranded DNA of ~6400 nt (Day et al., 1988;Marvin et al., 2014). The virion is composed of five proteins. The major coat protein pVIII, that forms the shaft of the filament, is present in the virion in a few thousand copies. Two pairs of minor coat proteins, pIII/pVI and pVII/pIX, form two asymmetric ends of the filament. They are each present in the virion in up to five copies. Infection with filamentous phage is mediated by binding of a minor protein, pIII, to the primary and secondary receptors, the tip of the F-pilus and the TolQRA protein complex spanning the periplasm and the inner membrane, respectively. The primary receptor, however, is not essential for infection; in the presence of Ca^2+^ ions as many as 1% of *E. coli* cells in the culture can be infected with Ff (Russel et al., 1988). TolQRA, a conserved protein complex in Gram-negative bacteria, appears to be both an essential and universally required protein for filamentous phage infection, allowing what appears to be a low-efficiency broad-spectrum infection of Gram-negative bacteria by this group of phage (Heilpern and Waldor, 2000). After entry into the host cytoplasm, the ssDNA genome [the positive (+) strand] replicates as an episome via a rolling-circle mechanism. First, the negative (-) strand is replicated by host proteins using the (+) strand template, forming the double-stranded circular supercoiled replicative form (RF) (Zenkin et al., 2006). The RF then serves as a template for (+) strand replication, which is initiated and terminated by the phage-encoded strand-transferase pII (Asano et al., 1999). The (+) strands accumulate in the cytoplasm in complex with phage-encoded ssDNA-binding protein pV, which covers the whole genome apart from a hairpin loop, called the packaging signal (PS). The ssDNA-pV complex is targeted to assembly sites at the inner membrane by PS. The (−) and (+) strand origins of replication, as well as the packaging signal, are together often referred to as the f1 origin of replication and are located within a ~400 nt long intergenic sequence (IG; Suppl. Fig. 1A). The (+) strand origin, or (+) origin, has a core (I) region and an extended (II) region. Both regions together form a binding site for the phage-encoded replication protein pII. In the absence of region II, the replication level from the (+) origin is about 1% that of the wild-type, but is increased to a nearly normal level by compensatory mutations (IRI) in *gII* encoding pII replication protein (Dotto et al., 1984). Region I also contains the pII nicking site that is absolutely required for initiation of (+) strand replication.

When inserted into plasmids, the intergenic sequence (f1 origin) allows rolling circle replication in the presence of helper phage, followed by packaging of the resulting ssDNA into Ff-derived particles; such vectors are known as phagemids. Some phagemid vectors are designed to express translational fusions with the Ff virion proteins and are used in phage display technology (Vieira and Messing, 1987;Barbas et al., 1991).

The virion proteins are all integrated into the inner membrane prior to assembly. The Ff virions are assembled by transfer of proteins from the inner membrane onto the ssDNA genome through a mechanistically and structurally poorly understood secretion-like process, mediated by phage-encoded trans-envelope secretion-assembly machinery (Rakonjac et al., 2011). Whereas the diameter of the wild-type Ff virion is constant, the length depends on the amount of packaged ssDNA. The shortest Ff-derived particles reported were 50 nm in length (Specthrie et al., 1992) and were produced by helper-phage-assisted replication and assembly of particles containing circular ssDNA of 221 nt. This short ssDNA was produced by replication from a specially constructed origin of replication, 303 nt in length (Suppl. Fig. 1B), inserted into a plasmid. This 303-nt sequence comprises two (+) origins of replication, *ori1* and *ori2*. The *ori1* serves as an initiator, but lacks region II. The *ori2* is further truncated and cannot bind pII or serve as an initiator; its role is to terminate replication of the (+) strand. In the presence of the phage-encoded replication protein pII, a segment between the two pII cut sites (TTCTTT↓AATA) in the two (+) origins is replicated to form a circular 221 nt ssDNA molecule. The two (+) origins flank a packaging signal required for assembly of this short circular ssDNA into 50 nm-long Ff-like particles, in the presence of a helper phage (Specthrie et al., 1992).

The physical properties of Ff phage, coupled with their amenability to genetic engineering using recombinant DNA technology, have enabled their extensive use in modern biotechnology and nanotechnology. Ff is central to phage display, a combinatorial technology in which libraries of peptides, antibodies or proteins are displayed on the virion surface, whilst the corresponding coding sequences are encapsulated inside the virions. This physical link between the displayed protein and its coding sequence allows affinity screening and enrichment of rare variants that bind to a ligand or a bait from vast libraries of variants (Smith, 1991;Rebar and Pabo, 1994;Zwick et al., 1998;Bradbury and Marks, 2004). The Ff phage have more recently been used as a nanoparticle-template to display arrays of organic and inorganic molecules (Bernard and Francis, 2014) for applications ranging from tissue targeting (Souza et al., 2010) and drug delivery (Bar et al., 2008) to nanoelectrodes (Lee et al., 2009), light-harvesting (Dang et al., 2013) and diagnostic devices (Petrenko, 2008). Furthermore, the liquid crystalline properties of Ff have been exploited to assemble colloidal membranes and other structures (Gibaud et al., 2012) and for applications in tissue engineering (Chung et al., 2011) and colorimetric sensors (Oh et al., 2014).

The current applications of Ff phage could be expanded by manipulating the length of the particles, potentially resulting in nanomaterials of novel properties. Short rods may be preferred over the long filaments in some applications, such as diagnostic methods that use diffusion (lateral flow) of diagnostic particles through complex matrices. Furthermore, short particles lacking viral or antibiotic-resistance genes would lower regulatory hurdles and consumer concerns, allowing wider application outside of laboratory containment.

To expand the versatility and decrease the risks Ff-derived nanoparticle use, we have developed a system for high-efficiency production of short functionalized Ff-derived particles (50 nm x 6 nm) that we named “Ff-nano”. These particles do not carry any genes and cannot replicate inside a bacterial cell. We show that these short particles are more resistant to heating in the presence of ionic detergent SDS compared to the full-length phage. Furthermore we demonstrate that functionalized Ff-nano can be used as detector particles for a high-sensitivity dipstick assay detecting a test analyte, fibronectin, at a concentration of 0.35 femtomoles/µL (2 × 10^8^ molecules/ µL).

## 2. Material and Methods

### 2.1. Bacteria, Phage and Growth Conditions

All *E. coli* strains used in this study are derivatives of laboratory strain K12. They were propagated in Difco™ 2xYT (Yeast Extract Tryptone) liquid medium (Becton-Dickinson (BD) and Company USA) at 37 °C with continuous shaking (200 rpm), or on 2xYT agar plates (1.2% Bacto-Agar, BD) at 37 °C unless otherwise stated. The medium was supplemented with suitable antibiotics where required, sourced from Sigma-Aldrich and Goldbio (USA). Antibiotics were used at the following concentrations: Ampicillin (Amp) 100 µg/mL, Kanamycin (Km) 30 µg/mL, Chloramphenicol (Cm) 25 µg/mL and Tetracycline (Tet) 15 µg/mL unless otherwise indicated.

Strain K561 (*HfrC λ^+^ relA1 spoT1 T2^R^ (ompF627 fadL701) ΔlacZ lac1^q^;* the Rockefeller University collection) was used for growth of the helper phage R408 (f1 *ΔPS gIX(T30A) IRI gtrxA2*); (Russel et al., 1986). Strains containing a *supD* mutation, K1030 [*HfrC λ^−^ relA1 spoT1 T2^R^ (ompF627 fadL701) supD zed508::Tn10;* the Rockefeller University collection) and K2092 (*supD, supE,* Δ*(hsdM-mcrB)5* (rk- mk- McrB-), *thi, Δ(lac-proAB), zed508::Tn10*, F′ [*traD36 lacI^q^ Δ(lacZ)M15 proA^+^B^+^*]; this work) were used for growth of *gVIII^amber^* mutant phage R777 (R408, *gVIII^am^25*) or Rnano3 (R777, MCS in *gIII)*. K2092 was obtained by co-transducing the *supD* mutation with the *zed508:*:Tn10 (Tet^R^ marker) from strain K1030 into strain TG1 *(supE, Δ(hsdM-mcrB)5* (rk- mk- McrB-), *thi, Δ(lac-proAB)*, F′ [*traD36 lacI^q^ Δ(lacZ)M15 proA^+^B^+^*] (Carter et al., 1985) using generalized transduction with P1 (Cm^r^ Clr-100) (Sternberg and Maurer, 1991). Strain K1030 containing pIV-expressing plasmid pPMR132 (Linderoth et al., 1997) was used for growth of phage R676 (f1, *ΔgIV, gVIII ^am^*; (Feng et al., 1999).

### 2.2. Construction of Phage and Plasmids

General molecular biology techniques for cloning, PCR amplifications and sequencing were carried out as described in (Sambrook and Russell, 2001). Molecular biology reagents were sourced from New England Biolabs Inc., (USA), Roche Molecular Biochemicals (Germany), Life Technologies Inc. (USA) and Takara (Japan). Oligonucleotides used in cloning, sequencing and PCR reactions were manufactured by Life Technologies Inc. and Integrated DNA Technologies Inc., (USA). DNA sequencing was carried out at the Massey University Genome Services, Institute of Fundamental Sciences, Palmerston North, New Zealand. The small and large scale plasmid and phage closed-circular double-stranded (RF) DNA was prepared using High Pure Plasmid Isolation Mini and Midiprep kits (Life Technologies; Roche Molecular Biochemicals) according to the manufacturers’ instructions.

Helper phage R777 was constructed from phage R408 and R676 (Feng et al., 1999), by combining the R408 origin of replication and *gII* (IRI) with *gVIII^am^* of R676. The RF (double-stranded) DNA of R408 and R676 was cut with *BamHI* and *BsrGI* The purified large *BamHI/BsrGI* fragment of R408 containing genes *gVI, gI, gXI, gIV, gII* and most of *gV* was ligated to the purified small *BamHI/BsrGI* fragment of R676 containing *gVII, gIX, gVIII* and of the 5’ moiety of *gIII*. The ligation was transformed into electrocompetent cells of the *supD* strain K1030. To identify R408 *gVIII^am^*, plaques obtained in the transformation plate were streaked onto the lawns of suppressor strain K1030 and a non-suppressor strain K561. Recombinant phage clones that formed plaques on K1030, but not K561 transformed with pIV producing plasmid pPMR132, were consistent with the correct recombination product (replication in the absence of pIV due to a wild-type *gene IV*, but requiring the *supD* mutation due to the presence of the *gVIII^am25^* mutation).

The helper/vector phage Rnano3 was constructed by combining the *gVIII^am25^* mutation with a 45-nt multiple cloning site (MCS) identical to that of the phage display vector pHEN2 (Marks et al., 1991) in the R408 background. A fragment containing both of these changes was amplified by overlap-extension PCR, using R777 as a template, flanking primers NJB6000 (5’-GTGCCTTCGT AGTGGCATTA-3’) and NJB6004 (5’-ACATAAATCA ATATATGTGA GTGA-3’) and overlapping primers containing multiple restrictions sites, NJB6001 (5’-GTTCCTTTCT ATTCTCACTC CGCGGCCCAG CCGGCCATGGG ATATCAGGCGGC-3’) and NJB6003 (5’-CCATGGGATA TCAGGCGGCC GCTCCCGGGG GCGCTGAAAC TGTTGAAAGT TGTT-5’). The product was designed to amplify the flanking *gVIII^am25^* allele and *gIII* sequences of R777, including the restriction sites *SnaB*I and *BamH*I. These sites were used for inserting the product into the *SnaB*I-*BamH*I-cut phage R408. The correct recombinants were identified by the requirement of a *supD* mutation in the host for plaque formation as described above, except that strain K2092 served as a host to avoid restriction of the PCR-amplified insert by the rk^+^, mk^+^ strain K1030. The resulting helper phage-display vector was named Rnano3. It contains a multiple cloning site identical to the pHEN2 vector, including unique restriction sites for enzymes *Sfi*I, *Sac*II, *Bgl*II, *Nco*I, *EcoR*V, *Xma*I, *and NotI*. Another helper/vector, named R408-3 and derived from the standard (*gVIII^+^*) helper phage R408, was constructed using the same strategy, except that the template for amplification was R408 (rather than R777).

To construct a recombinant helper/vector Rnano3FnB, displaying Fibronectin Binding Domain (FnB) from *Streptococcus pyogenes* (Rakonjac et al., 1995), the coding sequence for FnB was amplified from plasmid pDJ04 (Jankovic et al., 2007) using primers JR438 (5’-TCCCCCGCGG GAGGTCATGG ACCGATTGTC-3’) and JR439 (5’-TCCCCCCGGG CTCGTTATCA AAGTGGAAGA AGC-3’). The JR438 and JR439 primers introduced a *Sac*II site and an *Xma*I site, respectively, at the ends of the product, which was cleaved with corresponding restriction enzymes and inserted into a *Sac*II-*Xma*I-cleaved Rnano3. The ligation mixture was transformed into strain K2092. The correct recombinants were verified by restriction analysis and sequencing.

The Ff-nano-producing plasmid pNJB7 was constructed by inserting the Ff-nano origin of replication, amplified using plasmid pLS7 (Specthrie et al., 1992) as a template and primers NJB26 (5’-AGACGTTTTCCAGTTTGGAACAAG-3’) and NJB28 (5’-CCTATAAAAATAGGCGTATCACGAG-3’) into the vector pCR4-TOPO using the corresponding cloning kit (pCR4-TOPO blunt; Life Technologies, USA).

### 2.3. Phage Growth

All helper-vector phage stocks (R777, Rnano3 and Rnano3FnB) were prepared initially from a single plaque using a plate method. Briefly, phage were extracted from a plaque into 1 mL of 2xYT by slow rotation at room temperature for 1 h, filtered through a 0.45 µm-pore filter to remove the *E. coli* cells and titrated on an appropriate strain. To make a plate stock, 10^5^ - 10^7^ phage from the dissolved plaque were mixed with a culture of an appropriate host in 2xYT soft agar, plated and incubated overnight. The phage were extracted into 5 mL of 2xYT overlaid onto the surface of the lawn, followed by slow shaking for 1 h at room temperature. The 2xYT containing extracted phage was collected and filtered to remove *E. coli*. Titers of the plate stocks were typically ~10^11^ per mL. These plate stocks were then used for preparation of larger-volume stocks using a standard single-round infection of exponentially growing liquid cultures at a multiplicity of infection (m.o.i.) of 50 phage per bacterium.

### 2.4. Purification of the Ff-nano Particles

An exponentially growing culture (2 - 8 L) of K1030 carrying pNJB7 (Ff-nano-producing plasmid) was infected with appropriate helper phage (R777, Rnano3 or Rnano3FnB) at an m.o.i. of 50 and incubated at 37 °C without shaking for 20 min, to allow infection. Incubation was continued overnight at 37 °C with shaking at 200 rpm. The subsequent day, cells were pelleted (7000 x g for 20 min at 4 °C) and the supernatant containing the Ff-nano and full-length helper phage was subjected to differential two-step PolyEthylene Glycol (PEG) precipitation, to separate the two types of particles from each other, as described (Specthrie et al., 1992). Briefly, in the first precipitation, culture supernatant was subjected to 2.5 % w/v PEG8000 / 0.5 M NaCl and incubated overnight at 4 °C to precipitate the full-length helper phage. Precipitated full-length helper phage were collected by centrifugation at 16,500 x g for 45 min at 4 °C. Supernatant containing the Ff-nano particles was transferred to sterile containers, whereas the full-length phage pellet was suspended in TBS (50 mM Tris, 150 mM NaCl, pH 7.0). In the second precipitation, solid PEG8000 was added to the supernatant, to a final concentration of 15 %, and incubated overnight at 4 °C. The following day the Ff-nano-rich precipitate was collected by centrifugation (at 16,500 x g for 45 min at 4 °C) and the pellet was resuspended in TBS. The full-length phage and Ff-nano fractions (resuspended pellets from 2.5 % and 15 % PEG, respectively) were further subjected to successive detergent treatments and precipitations to decrease contamination with fragments of inner and outer membranes [Sarkosyl (1 % w/v) and Triton X-100 (0.1 % v/v)]. Full length helper phage was recovered after each detergent treatment by precipitation with 2.5 % PEG and Ff-nano with 15% PEG as described above, except that precipitations were carried out at room temperature to decrease co-precipitation of detergents with the particles.

Ff-nano particles were further purified away from the remaining full-length helper phage based on the large difference in size. Native agarose gel electrophoresis was carried out as described (Nelson et al., 1981;Rakonjac and Model, 1998). Preparative native agarose gel electrophoresis was carried out on 0.8 % agarose gels. Upon completion of electrophoresis, the agarose gel slabs containing the bands corresponding to full-length or Ff-nano particles were separately cut out and transferred to dialysis tubes (Novagen, D-tube dialyzer Maxi, MWCO 12–14 kDa) containing 500 μL of sterile 1 × TAE buffer (40 mM Tris-acetate buffer, pH 8.3, 1 mM EDTA). These tubes were then placed into an electrophoresis chamber in sterile 1 × TAE buffer and electrophoresed overnight at 0.5 V/cm at 4 °C. After removing the gel slices from tubes, the phage particles (full-length or Ff-nano) were precipitated overnight at 4 °C using 2.5 % or 15 % PEG, respectively, in 0.5 M NaCl. The precipitate was dissolved in an appropriate amount of 1 × PBS (137 mM NaCl, 2.7 mm KCl, 10 mM Na_2_HPO_4_, 1.8 mM KH_2_PO_4_ and stored at −80 °C until further use.

### 2.5. Quantification of Ff-derived Particles

Phage R777, Rnano3 and Rnano3FnB (containing an amber mutation in gVIII) were titrated on *E. coli supD* strains K1030 or K2092 to determine the number of plaque-forming units per milliliter, using standard phage plating and titration methods. To quantify non-infectious Ff-nano, the particles were disassembled in SDS-containing buffer (1 % SDS, 1 × TAE, 5 % glycerol, 0.25 % Brom-Phenol Blue) at 100 °C for 5 min and subjected to electrophoresis in 1.2 % agarose gel in 1 × TAE buffer. After electrophoresis, the ssDNA released from disassembled Ff-derived particles was stained with ethidium bromide (Suppl. Fig. 2) and quantified by densitometry (Rakonjac and Model, 1998). The amount of ssDNA in any band is not linearly proportional to the intensity of fluorescence, therefore each gel contained a set of twofold dilutions of purified full length f1 ssDNA that were used for calibration (Suppl. Fig. 2). Gels were photographed by a CCD camera (Biorad, USA). Image Gauge V4.0 and Microsoft Excel were used for densitometry and calculations, respectively. A second-order polynomial function was used to fit the standard curve. Amounts of ssDNA were converted to the genome equivalents based on the molecular weight of the ssDNA genome. The molecular weight of the Ff-nano ssDNA genome was in turn calculated from its length (221 nt) and nucleotide composition (derived from the DNA sequence).

Native agarose gel electrophoresis was used to separate the intact full-length (helper) phage from Ff-nano particles, for analysis and purification. The bands containing intact particles were detected in agarose gels after the in situ stripping the virion proteins off the ssDNA by soaking the gel in 0.2 M NaOH for 1 h, followed by neutralization with 0.45 M Tris pH 7.0 and staining with ethidium bromide (Suppl. Fig. 3).

### 2.6. Microscopy

The phage samples for non-cryo TEM analysis were adsorbed onto glow-discharged carbon-coated grids, negatively stained with a 1% uranyl acetate solution and examined on a Phillips CM-10 microscope at the Manawatu Microscopy and Imaging Centre (MMIC, Institute of Fundamental Sciences, Massey University).

Cryo-negative staining was performed at the University of Pittsburgh, PA, USA, on an FEI (Hillsboro, OR) Tecnai F20 microscope equipped with a Gatan (Pleasanton, CA) 626 cryoholder and operated at 200 kV and nominal magnification of 50,000×. Briefly, 3 µl of sample were pipetted onto a holey carbon/copper grid that had been briefly glow-discharged. The grid was then placed sample-down onto a 100 µl droplet of 16 % ammonium molybdate (in the pH range 7.0 to 8.0) and floated for 60 sec, following the cryo-negative staining procedure (Adrian et al., 1998). The grid was then removed, blotted and plunge-frozen into liquid ethane using an FEI Vitrobot Mk III. Images were collected using standard low-dose techniques on a Gatan Ultrascan 4000 CCD camera with post-column magnification of 1.33×, yielding a pixel size at the sample of 2.3 Å.

### 2.7. Phage ELISA Assay

Phage Enzyme Linked Immunosorbent Assay (ELISA (Harlow and Lane, 1999)) was carried out in the 96-well plates (Nunc-Immuno MaxySorpTM, Denmark). The wells were coated overnight at 4 °C with 100 µL of human plasma fibronectin (Sigma-Aldrich, USA) at 20 µg/mL in 100 mM sodium bicarbonate buffer, pH 9.5. All subsequent steps were carried out at room temperature. The wells were washed once with 300 µL of PBS containing 0.05 % Tween 20 (PBST) and blocked with 250 µL of 2 % (w/v) Bovine Serum Albumin (BSA) in PBS for 2 h. After washing wells three times with 300 µL of PBST, Rnano3FnB full-length phage or Ff-nano particles (1 × 10^8^) in 100 µL of PBS were added to the wells. Negative controls (buffer and protein) were PBS and 2% BSA in PBS, respectively, whereas negative phage controls were the full-length phage and the Ff-nano particles derived from “empty” vector/helper Rnano3 (and therefore not displaying FnB). The plates were then incubated for 2 h. The unbound particles were removed by extensive washing with PBST (300 µL, 7 times). To detect phage particles bound to fibronectin in the wells, 100 µL of rabbit anti-pVIII (polyclonal antibody to M13, fd and f1, Progen Biotechnik; Germany) was added at the concentration of 0.1 µg/mL in 1 × PBS and incubated for 1 h. The wells were then washed five times with 300 µL PBST buffer. Next, 100 µL of secondary HRP-conjugated anti rabbit antibody was added to all wells at a dilution of 1:2000 and then washed with 300 µL PBST seven times. The HRP bound to the plate was detected with 1-StepTM Turbo TMB-ELISA reagent (Thermo Scientific). The enzyme reaction was stopped by adding 25 µL of 0.5 M H_2_SO_4_. The signal was quantified by measuring absorbance at 450 nm.

### 2.8. Labelling of Ff-derived Particles Using Fluorescein Isothiocyanate

Fresh Fluorescein IsoThioCyanate (FITC) solution (1 mg/ml) was prepared in 1 M NaCO_3_ / NaHCO_3_ (pH 9.0) buffer. Ff-nano (0.5 × 10^13^ in 500 µL) was precipitated using 15 % PEG / 0.5 M NaCl and the same amount of the full-length phage with 2.5 % PEG / 0.5 M NaCl. The precipitate was dissolved in 200 µl of 1 M NaCO_3_ / NaHCO_3_ (pH 9.0) buffer. The FITC solution (50 µL) was added to the suspension of precipitated Ff-nano or full-length phage and the reaction mixture was rotated in the dark for 1 h at room temperature. The reaction was stopped by adding 10 µl of the NH_4_Cl and the particles were purified by a series of two PEG8000 precipitations using the concentrations appropriate for Ff-nano (15 %) and full-length phage (2.5 %). The pellet obtained after the second PEG8000 precipitation was suspended in 100 µl of 1 × PBS and stored at 4 ºC in dark until further use.

### 2.9. Dipstick Assays

Hi-Flow Plus membrane cards were used to make the dipsticks (Millipore Corporation). These cards contain nitrocellulose membranes cast on a 2 mil (0.001 inch; USA) polyester film backing. The reagents for the test and control lines were applied to the membrane using a mechanical dispenser. Collagen I (Sigma; 1 µg/µL in 0.25 % acetic acid;) was dispensed at the test line, whereas undiluted mouse anti-pVIII antibody or anti-FITC antibody (0.5 to 1 µg/µL) was dispensed at the control line. The protein-loaded cards were allowed to dry at 37 °C for 1 h and were subsequently cut into 0.5 cm × 2.5 cm size dipsticks using an automatic card-cutting tool (BIODOT membrane cutter; SM5000; sheet slitter). The dipsticks were stored in zip-lock sealed bags at room temperature in a dark place until use. The membrane has a flow rate of 46 sec / 2.5 cm (total length of the dipstick).

The distal end of protein-loaded dipstick was dipped into the 50 µL mixture of analyte (fibronectin, 5 µg/50 µL for initial detection and serial two fold dilutions starting from 0.5 µg/50 µL for quantitative detection) and the detection particles. Full length phage were used at 1 × 10^10^ per assay, and Ff-nano (or Ff-nanoFnB) particles at 1 × 10^11^ per assay. Dipsticks were allowed to stand in solution for 30 min and dried for 1 h at 37 °C. The FITC-labelled particles on the dipsticks were detected using a phosphoimager (Fuji, Japan). Unlabeled particles were visualized by western blotting using rabbit pVIII-specific antibody (0.66 μg/mL; Progen Biotech) for 1 h at room temperature, followed by 5 washes of 5 min each with PBST. Rabbit IgG-specific antibody conjugated to alkaline phosphatase (Sigma, USA) was used as the secondary antibody. The dipsticks were washed again five times with PBST, and developed using Nitro Blue Tetrazolium (NBT) and 5-Bromo-4-Chloro-3-Indolyl Phosphatase (BCIP) in alkaline buffer (Blake et al., 1984).

## 3. Results

### 3.1. The Ff-nano Production and Purification System

In this work we have constructed an Ff-derived phage display system for production of functionalized nanorods (50 x 6 nm) we named Ff-nano (Fig. 1). In the first stage, we modified a setup for production of short phage originally published by Specthrie *et al*. (1992) to increase the amount (per cell) of the 221-nt circular ssDNA available as a template for assembly of the short particles. This was achieved by inserting the 303-nt short-phage (microphage or Ff-nano) origin of replication (Suppl. Fig. 1) into the high-copy-number plasmid pCR4-TOPO (in contrast to the low-copy-number plasmid pBR322 used in the original system), to obtain plasmid pNJB7. Furthermore, a new helper phage, named Rnano3, was constructed. The helper phage Rnano3, in contrast to the original helper phage R474 used for short phage production (Specthrie et al., 1992), is not only a helper but also a phage display vector. Rnano3 was designed for display of proteins as fusions with phage protein pIII that is present in up to 5 copies at one end of the Ff-derived particles. Infection of cells containing plasmid pNJB7 with the helper phage/vector Rnano3 resulted in production of the 50 nm x 6 nm particles (Ff-nano or microphage; Fig. 2) and full-length phage.

**Figure 1.**
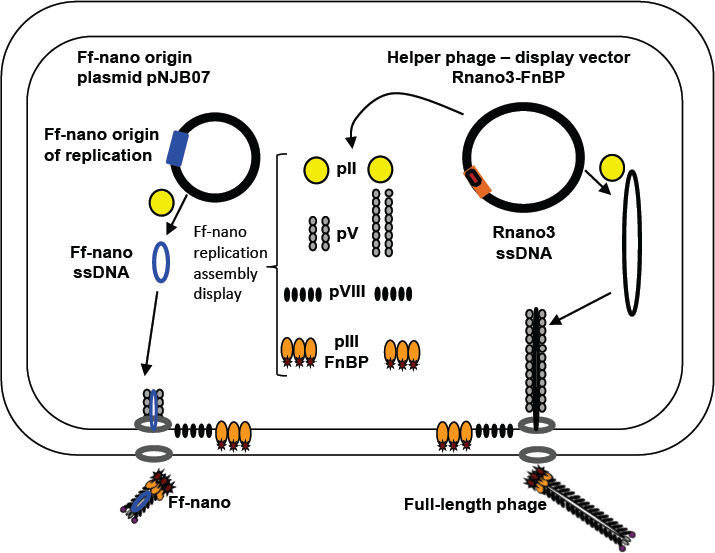
The system for production of functionalized Ff-nano. *E. coli* cells containing the Ff-nano production plasmid pNJB7 were infected with the helper phage Rnano3FnB containing the coding sequence of a “probe” or “detector” protein fused to pIII. Upon infection, pII from the helper phage induces positive strand replication from the pNJB7 Ff-nano origin of replication and also provides all other phage proteins and assembly machinery for production of the Ff-nano particles. All five copies of pIII are fusions to the probe (only three copies of pIII fusions are shown).

The 50-nm-particle production was originally reported to be improved in the presence of helper phage that produces decreased amount of the major coat protein pVIII. Decreased amount of pVIII presumably favors initiation and termination of assembly over elongation of particles, thereby decreasing the co-packaging of short 221 nt ssDNA with the helper phage DNA into the full-length particles and increasing production of short particles (Specthrie et al., 1992). In contrast to R474 helper used previously for the short phage production, which has a mutation lowering transcription of not only *gVIII* but also in *gIX* (Russel and Model, 1989;Specthrie et al., 1992), the Rnano3 helper phage/vector that we constructed produces decreased amount of pVIII without affecting *gIX* expression. This specific effect on pVIII is achieved by using a suppressed *gVIII^am^* mutation, *gVIII^am25^*, which contains a TAG stop codon in the *gVIII* sequence at codon 25 (replacing the GAG codon for Glu). When the Rnano3 phage infects a *supD* strain, the TAG codon is translated into Glu or Ser (Rogers and Soll, 1988). The efficiency of the *gVIII^am25^* mutant translation in the *supD* strains (K1030 and K2092) is <50% that of the wild-type, resulting in a lower number of pVIII copies produced per cell than in an infection with phage containing a wild-type *gVIII*. Besides the helper/vector with decreased amount of pVIII, we also constructed and examined helper/vector R408-3 that is identical to Rnano3 apart from having a wild-type *gVIII*. Rnano3 and R408-3 were both found to support production of the Ff-nano particles (Suppl. Fig. 4). As predicted, co-packaging of the 221-nt ssDNA derived from the Ff-nano origin of replication with the full-length phage genome appeared to be increased in the R408-3 full-length phage fraction after differential PEG precipitation, relative to that of Rnano3 (*gVIII^am25^* mutant), however the production of Ff-nano particles was also increased (Suppl. Fig. 4).

Ff-nano particles were partially purified from the helper phage by differential PEG precipitation (see Material and Methods section). The full-length helper phage was first precipitated out of the culture supernatant in 2.5 % PEG8000, 0.5 M NaCl, followed by increase in PEG8000 to 15 % to precipitate Ff-nano (Specthrie et al., 1992). However, this method resulted in 0.7 % full-length phage still remaining in the Ff-nano fraction (Table 1). We used native agarose gel electrophoresis to separate the short particles from the full-length phage following the differential PEG precipitation (Suppl. Fig. 3). Preparative agarose gel electrophoresis is simple, fast, effective and less expensive than the size fractionation by Sepharose CL2B columns used by Specthrie et al. (1992). The band containing Ff-nano was excised and the particles were extracted by electroelution (see Material and Methods for details of purification). The native preparative agarose gel electrophoresis purification step decreased the full-length helper phage frequency in the final purified sample by a factor of 1400, down to 5.0 × 10^−6^ relative to Ff-nano (Table 1). This method was also relatively efficient in recovery of the Ff-nano (31% of the input; Table 1).

**Table 1.** Purification of the Ff-nano by native agarose gel electrophoresis and electroelution.

|  | **Ff-nano**<br>**(total**<br>**number<sup>c</sup>)** | **Recovery**<br>**of the Ff-**<br>**nano<sup>d</sup>** | **Helper**<br>**phage**<br>**(total**<br>**number<sup>e</sup>)** | **Helper : Ff-**<br>**nano ratio<sup>f</sup>** | **Fold decrease of**<br>**helper : Ff nano**<br>**ratio<sup>g</sup>** |
| --- | --- | --- | --- | --- | --- |
| **Input<sup>a</sup>** | $6.4 \times 10^{14}$ | N/A (input) | $4.5 \times 10^{12}$ | $7.0 \times 10^{-3}$ | N/A (input) |
| **Electro-**<br>**purified**<br>**Ff-nano<sup>b</sup>** | $2.0 \times 10^{14}$ | 31 % | $9.9 \times 10^8$ | $5.0 \times 10^{-6}$ | 1400 |

<sup>a</sup> Starting material loaded onto the preparative gel corresponds to the Ff-nano-enriched lysate obtained by differential PEG precipitation.
<sup>b</sup> Ff-nano-band was excised from the agarose gel, extracted by electroelution, purified and concentrated (see Material and Methods section for details).
<sup>c</sup> Determined by densitometry (see Material and Methods section for details).
<sup>d</sup> Ratio of the Ff-nano number in the electro-purified sample to the amount in the input.
<sup>e</sup> Determined by titration.
<sup>f</sup> Determined by dividing the total number of full-length helper phage with the total number of the Ff-nano particles.
<sup>g</sup> Fold decrease of the helper to Ff-nano ratio (“helper : Ff-nano ratio”) was calculated by dividing the “helper : Ff-nano ratio” in the input with the “helper : Ff-nano ratio” in the electro-purified sample (data column four; row one divided by the row two value).

### 3.2. Physical Properties of the Ff-nano

#### 3.2.1. Morphology

Purified Ff-nano were negatively stained and visualized by cryo-electron and transmission electron microscopy (Fig. 2). Dimensions of the Ff-nano are 50 nm by 6 nm, matching those reported in Specthrie et al. (1992). The Ff-nano particle termini appear asymmetric, one pointy and one blunt, as reported for the Ff phage (Gray et al., 1979). In the cryo-negative micrographs Ff-nano formed sheets composed of individual Ff-nano aligned in alternating orientations. In addition, in a larger field, about 3.5 % (6 out of 172) double-length Ff-nano particles could be observed (Suppl. Fig. 5). This is consistent with observations of full-length filamentous phage, which, depending on the genotype, contain some proportion of particles that are longer than the majority by a factor of two or multiple lengths of the virion, and containing two or more sequentially packaged genomes (Rakonjac and Model, 1998). We note that the Ff-nano particles visualized in the electron micrographs do not have any signs of the extra balls of density that are sometimes observed attached to the pIII end of the filament and correspond to free-moving N1N2 domains of pIII (Gray et al., 1979).

**Figure 2.**
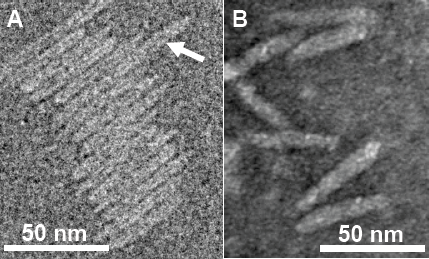
Transmission electron micrographs (TEM) of Ff-nano particles. (**A**) Cryo-negative TEM of particles obtained using the R777 helper phage. (**B**) Negative-stain TEM of particles obtained using the Rnano3 helper/vector. Arrow in (A) points to a double-length particle. Samples were prepared as described in Material and Methods.

#### 3.2.2. Stability to Heating in the Presence of Sodium Dodecyl Sulfate

In the course of Ff-nano analyses, we observed that the standard protocol for Ff phage *in vitro* disassembly, heating in a buffer containing 1 % (34 mM) ionic detergent sodium dodecyl sulfate (SDS) for 5 min at 70 °C, was not efficient in releasing ssDNA from the Ff-nano particles (data not shown). This indicated that the Ff-nano particles could be more stable to heat/SDS treatment than the full length helper phage, even though both are assembled within the same *E. coli* cell. To test this hypothesis, a time-course experiment of heat exposure was used to monitor disassembly of full-length helper phage and Ff-nano isolated from the same culture using differential PEG precipitation and preparative agarose electrophoresis, as described in the previous section. Approximately 2 × 10^12^ Ff-nano or 1 × 10^11^ full-length phage were heated at 70 °C in the presence of 1 % SDS for a period from 5 to 20 min; one sample was also incubated at 100 °C for 5 min. Disassembly of the helper (full-length) phage and Ff-nano particles was monitored by agarose gel electrophoresis (Fig. 3). Released ssDNA was directly visualized by staining with ethidium bromide (Fig. 3B), while the ssDNA that remained encapsidated inside the intact Ff-derived particles (resistant to heat/SDS treatment) was not detectable by direct staining. In order to visualize the SDS-resistant intact particle bands after electrophoresis, the coat proteins were dissociated from ssDNA *in situ* by soaking the gel in an alkaline buffer (NaOH), followed by neutralization and re-staining of the gel by ethidium bromide (Fig. 3A).

**Figure 3.**
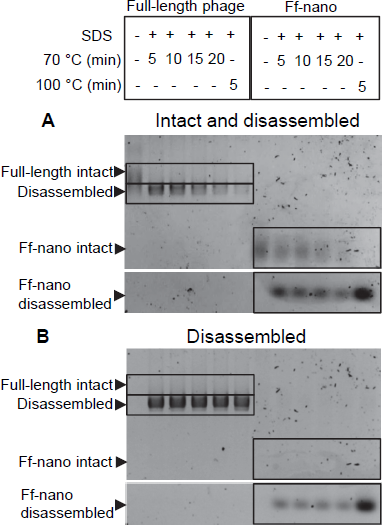
Ff-nano resistance to heating in SDS. (**A**) Intact (undamaged) particles and free ssDNA released from particles by the heat/SDS-treatment. Free ssDNA was visualized by staining the gel with ethidium bromide. Bands corresponding to the intact particles were visualized after soaking the gel in 0.4 NaOH (see Material and Methods for details). Gel sections containing the bands corresponding to the free ssDNA and intact full-length and Ff-nano particles are boxed. (**B**) Free ssDNA only, released by the heat/SDS treatment (visualized by direct ethidium bromide staining of the gel prior to NaOH treatment).

When the full-length (helper) phage virions were analyzed, the untreated samples (without SDS or heating) did not contain any free ssDNA. All full-length phage were therefore intact (all ssDNA was contained within the virion; Fig. 3; compare the corresponding untreated sample lanes in gels A and B). In the presence of 1 % SDS, at the first time-point (5 min) of incubation at 70 °C, however, all ssDNA was in the free form and none in the intact phage particles, hence the vast majority of the full-length phage particles were sensitive to these conditions (Fig. 3A). In contrast, when the Ff-nano particles were subjected to the same treatment (1 % SDS at 70 °C), a large proportion of the intact Ff-nano particles were detected, with some free ssDNA observed at all time-points (Fig. 3, compare the 70 °C-treated sample lanes in A vs. B). The amount of intact Ff-nano particles decreased gradually at 70 °C between the 5 min and 20 min time points, but nearly half stayed intact throughout the incubation. The intact Ff-nano were completely eliminated only after incubation at 100 °C, confirming that only at this higher temperature is SDS able to disassemble all Ff-nano particles in the sample (Fig. 3, compare lane 12 in A vs. B). This experiment demonstrates that the Ff-nano particles have superior resistance to heating in the presence of ionic detergent SDS in comparison to the full-length phage, requiring higher temperature (100 vs. 70 °C) for dissociation of most particles.

### 3.3. Functionalization of the Ff-nano

To test the potential of the Ff-nano-production system to assemble functionalized particles that can be used for display of foreign proteins, a fusion to pIII was constructed in the helper/vector Rnano3. The displayed peptide was the Fibronectin-Binding (FnB) domain from Serum opacity factor (Sof), a surface protein of *Streptococcus pyogenes* (Rakonjac et al., 1995). The Sof FnB domain is composed of 3 repeats of a bacterial Fibronectin-binding motif (PF02986) that each binds to the N-terminal domain of Fibronectin (Fn) with a low nanomolar dissociation constant (Rakonjac et al., 1995;Schwarz-Linek et al., 2006). The high affinity of FnB for the Fn makes it a good candidate as a detector molecule and FnB-Fn combination is a good detector-analyte pair to investigate the use of Ff-nano as a display particle and for its applicability to lateral flow dipstick diagnostic devices.

The functionalized helper/vector encoding the FnB-gIII fusion was named Rnano3FnB. When *E. coli* containing the Ff-nano-origin plasmid (pNJB7) was infected with Rnano3FnB, the Ff-nano particles displaying FnB (named Ff-nanoFnB) were produced together with the full-length helper phage (Fig. 1). The helper (full-length) Rnano3FnB phage and the Ff-nanoFnB particles obtained using this helper were separated by differential PEG precipitation and each of the long and the short phage were further purified using native agarose gel electrophoresis followed by electroelution, as described in the previous section. Phage ELISA was performed to confirm that the FnB domain was displayed (schematically represented in Fig. 4A). Rnano3FnB full-length and Ff-nanoFnB particles exhibited binding to immobilized fibronectin as indicated by strong ELISA signal detected for fibronectin-coated wells incubated with these phage particles (OD_450_ = 1.2 with 10^8^ particles per well in the presence of 40 ng/µL of Fn). No signal over background levels (OD_450_ = 0.1; intrinsic to the phage ELISA) was detected in control wells incubated with Ff-nano particles that did not display FnB, derived from the infection with vector Rnano3 (without the FnB domain) nor in the wells coated with BSA only (from which fibronectin was omitted; Fig. 4B), apart from the full-length Rnano3 particles (not displaying FnB) which gave a low signal OD_450_ = 0.3 in the presence of Fn. In conclusion, Ff-nanoFnB particles detected Fn and were hence displaying FnB. All Ff-nano and Ff-nanoFnB samples used in the assays were imaged by TEM to confirm the purity and morphology of the particles (data not shown).

**Figure 4.**
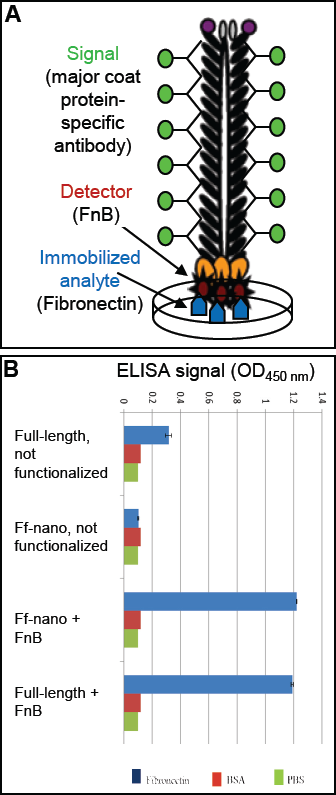
Confirmation of Ff-nano-FnB display by phage ELISA assay. (**A**) Schematic representation of the fibronectin-binding phage ELISA assay. (**B**) Assay result. Fibronectin was immobilized on a microtiter plate at a saturating concentration of 40 ng/μL (2 μg per well of a 96-well plate), whereas control wells for each assay were coated with PBS alone or BSA (1 %) in PBS. The wells were exposed to (1 × 10^8^) Ff-nanoFnB or full-length phage Rnano3FnB that both displayed FnB as a pIII fusion, or control particles Ff-nano and Rnano3 that did not display FnB. After washing of the wells, bound Ff-nano or full-length phage were detected using a primary antibody to the major coat protein, then the secondary HRP-conjugated antibody, followed by detection of HRP through an enzymatic reaction according to the standard ELISA protocols. Data are presented as an average of three measurements. Error bars show standard deviation.

### 3.4. Development of Quantitative assay for Fibronectin Using Ff-nano

#### 3.4.1. Dipstick Assays Using Ff-nano

To investigate the detection of fibronectin (the analyte) in solution using the Ff-nano displaying FnB domains as the detector (probe) in lateral flow devices, a simple dipstick assay was designed and tested (see Fig. 5A for a schematic representation of the Fn-detection dipstick assay). The dipsticks used in this assay contained human type I collagen at the test line (T). Collagen binds fibronectin with high affinity (Engvall et al., 1981).

**Figure 5.**
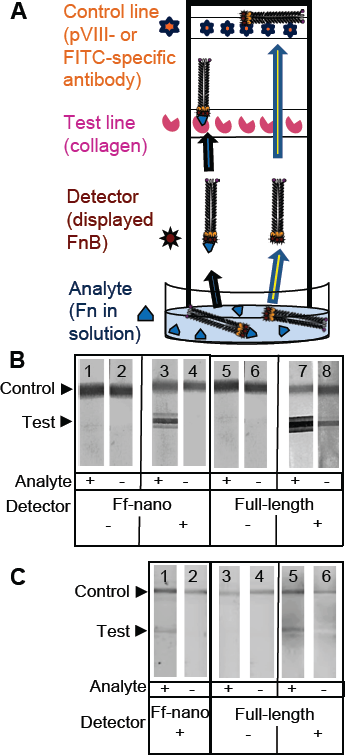
Dipstick assay using Ff-nanoFnB. (**A**) Schematic representation of a lateral-flow dipstick assay; Fn-detection dipstick assay using: (**B**) unlabeled; (**C**) FITC-labelled particles. Each assay (50 μL) contained 1 × 10^10^ full-length Rnano3 (Rnano3FnB) or 1 × 10^11^ Ff-nano (or Ff-nanoFnB) particles and 1 μg of Fn. The assay was performed and the unlabeled or FITC-labeled particles were detected as described in the Material and Methods section. The test line, printed with collagen solution, appears as a triple band when the signal is high. This is due to secondary lines flanking the main line that form during printing of collagen on the card. The triple banding during printing is caused by the acidity of the solution, necessary to keep the collagen soluble (0.25 % acetic acid).

The assays were carried out using particles that were either unlabeled (Fig. 5B), or FITC-conjugated (Fig. 5C). The unlabeled particles were captured on the control line using mouse pVIII-specific antibodies and detected on the dipstick, after the assay was completed, using rabbit pVIII-specific antibody. The FITC-labelled phage were captured on the control line using FITC-specific antibodies and detected on the dipstick using a phosphoimager.

As expected, the Ff-nanoFnB particles and full-length phage displaying FnB (Rnano3FnB) showed binding to collagen at the test line in the presence of fibronectin [Fig. 5B (sticks 3 and 7); Fig. 5C (sticks 1 and 5)]. The Ff-nanoFnB particles gave no background in the absence of Fn (analyte), whereas the full-length FnB-displaying phage showed some unspecific binding to the collagen test line (Fig. 5B, dipstick 8 and Fig. 5C, dipstick 6) in the absence of Fn. The Ff-nano and full-length particles that did not display FnB did not bind to the test (T) collagen line, indicating the lack of unspecific binding of phage to collagen [Fig. 5B (sticks 1, 2, 5 and 6); Fig. 5C (sticks 3, 4)].

#### 3.4.2. Quantification of Fibronectin Using Dipstick Assay

To examine the quantitative range and the limit of analyte detection using the Ff-nano dipstick assay, serial 2-fold dilutions of analyte (fibronectin) starting from 10 ng/µL were assayed using this format. Since the dipstick assay signal was stronger in the unlabeled particle assay (Fig. 5B) in comparison to the assay with FITC-labelled particles (Fig. 5C), the former were used for quantitative dipstick assays.

The intensity of the signal at the test line positively correlated with the concentration of analyte (Fig. 6A). Densitometric analysis of the signal at the test line (Fig. 6B) indicated a second order polynomial dependence between the signal intensity at the test line and analyte concentration over a range of concentrations between 1.25 ng/μL and 10 ng/μL. Concentration of Fn in the serum of healthy individuals is 259-400 ng/μL (Allard et al., 1986), Furthermore, variations of Fn concentration in disease are also within the range of quantitative detection of this assay (Choate and Mosher, 1983;Cembrowski and Mosher, 1984;Weller et al., 1988;Honest et al., 2002;Mosher, 2006;Eissa et al., 2010). Hence, suitably diluted samples of healthy and affected individuals can be quantitatively determined using this assay.

**Figure 6.**
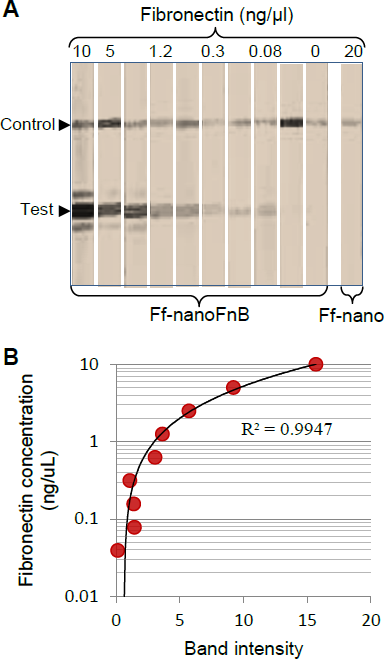
Quantitative fibronectin detection using Ff-nanoFnB. (**A**) Dipsticks exposed to decreasing concentrations of fibronectin; (**B**) Densitometry plot of the test line *vs*. Fn concentration. Unlabeled detection Ff-nanoFnB particles (2 × 10^11^) were mixed with Fn solution at the indicated final concentrations in a total volume of 50 µL for 30 min at room temperature. The assay was performed and the signal quantified as described in the Material and Methods section.

The lowest concentration of fibronectin that the assay could detect was 0.08 ng/μL, equivalent to 0.35 femtomoles/μL or 2 × 10^8^ molecules/μL, hence this is the sensitivity limit of the Ff-nanoFnB-based fibronectin detection dipstick assay that we developed.

## 4. Discussion

### 4.1. Ff-nano Production and Purification

Ff-derived functionalized nanoparticles reported to date are based on the long and thin filamentous template, whose hydrodynamic properties are not favorable for diffusion-based applications. From ethical and regulatory perspectives, the use of Ff-phage outside of the laboratory containment, in particular in home-use diagnostic devices or as carriers for tissue targeting of drugs, antigens or diagnostic markers, is potentially controversial because they are recombinant viruses. Due to the propensity of Ff to infect Gram-negative bacteria via the ubiquitous TolQRA complex in the absence of their primary receptor, the F-pilus of *E. coli* (Russel et al., 1988;Heilpern and Waldor, 2000), they could potentially infect a wide range of Gram-negative bacteria. This includes a possibility of mobilization and horizontal gene transfer of antibiotic-resistance-encoding genes, present in most of the phage display vectors, among the gut or environmental bacteria. The Ff-derived particles (virions) containing phage display vectors have been shown to be taken up by mammalian cells in culture (Burg et al., 2002), hence there is a possibility that the encapsidated DNA permanently inserts into patient genomes.

By functionalizing short Ff-derived particles that have no coding sequences and can therefore be considered a protein-DNA complex rather than recombinant virus, we have improved the prospect of Ff filamentous phage applications outside of the laboratory. Furthermore, the length-to-diameter ratio of these particles is 15-20-fold smaller than that of the Ff particles derived from standard phagemid and phage vectors, improving the diffusion rate of the particles.

We have developed a production and purification pipeline for 50 nm particles by converting an Ff phage short-particle production system (Specthrie et al., 1992) into a high-efficiency system for producing functionalizable nanoparticles that we named Ff-nano. The increase in efficiency was achieved by using a high-copy-number plasmid containing the Ff-nano origin of replication and construction of helper phage Rnano3 and R408-3 that are also phage display vectors, containing cloning sites for construction of protein fusions to pIII.

In the Ff-nano purification, we introduced size-separation from the full-length helper phage by preparative agarose electrophoresis and electroelution. This strategy achieved excellent resolution, leading to a high enrichment of the Ff-nano particles over the full-length helper phage at a low cost and high recovery of the Ff-nano particles (31%). Overall, this production and purification system yields around 10^14^ - 10^15^ Ff-nano particles from 2 - 8 L of *E. coli culture*. The Ff-nano samples obtained, however, still contain full-length phage at an approximate frequency of 1 in 200,000 (5 × 10^−6^). Although the helper phage that we use do not contain antibiotic resistance genes, they still represent an undesired population that needs to be eliminated in order to obtain fully virus-free Ff-nano samples.

### 4.2. Ff-nano Stability

The Ff-nano were more resistant to heating in 1% (34 mM) ionic detergent Sodium Dodecyl-Sulfate (SDS) in comparison to the full-length phage. This property is solely due to the difference in phage length, as the assayed populations of full-length and Ff-nano particles were purified from the same *E. coli* culture and therefore had identical protein composition. Ff filamentous phage dissociation in the presence of SDS was investigated in detail by Stopar and colleagues using Electron Spin Resonance (ESR), circular dichroism and NMR (Stopar et al., 1998; 2002; 2003). Disassembly of Ff by SDS is interesting as a model for dissociation of phage particles and insertion of pVIII into the lipid bilayer during phage infection. The process of virion dissociation is a transition between two states: i) a sub-solubilization state where phage particles are saturated with detergent molecules, but still coexist with the detergent in solution; ii) the solubilized state where coat protein is solubilized into detergent micelles (Stopar et al., 2003). These reports concluded that, in the sub-solubilization state, the detergent molecules become inserted or wedged into the grooves formed between pVIII monomers, thereby disrupting the hydrophobic protein-protein interactions of neighboring pVIII molecules. Solubilization of the major coat protein with detergent is an equilibrium of pVIII-pVIII interactions *vs*. pVIII-detergent interactions (Stopar et al., 1998). When a critical ratio of detergent to phage is reached, even a single pVIII subunit displacement across the filament triggers a steric change. This increases the entropy of the surrounding pVIII molecules due to the exposure of their hydrophobic surfaces to the solvent resulting in cooperative dissociation of the virion (Ikehara et al., 1975;Stopar et al., 2002). In our experiment, the full-length virions are rapidly and completely disassembled after incubation in 1 % SDS at 70 °C for 5 min, which is the time point when over half of the Ff-nano particles in the sample remain intact (Fig. 3). This length-dependent difference can be reconciled by the Ff-nano presenting a 17-fold smaller surface area for detergent interaction in comparison to the full-length phage, thereby decreasing the odds of steric imbalance caused by SDS-pVIII interactions per virion and resulting in much larger numbers of resistant particles under the same conditions. In addition, due to the short length of Ff-nano, the potential imperfections of pVIII packing that occur in each particle due to mechanical bending and twisting of the filament are greatly reduced. Bending can be observed in the electron microscopic images of full-length phage (Gray et al., 1979;Rakonjac and Model, 1998), but not Ff-nano (Fig. 2A; (Bennett, 2010). The short length, preventing the bending of the filament may therefore confer additional stability to Ff-nano in comparison to full-length virions.

From the technological standpoint, the increased stability of Ff-nano particles may be beneficial in applications that involve high temperature in detergent-containing environment, or other harsh conditions. Given the higher resistance of Ff-nano to heating in 1% SDS at 70 °C in comparison to complete sensitivity of the full-length phage, it is possible to develop conditions that will completely degrade the full-length phage in the purified Ff-nano preparation, while completely preserving the Ff-nano particles, thereby eliminating the full-length phage from the Ff-nano preparations. This approach will be possible for applications where the residue of SDS in the sample is not an issue, and where the Ff-nano particles display peptides that are resistant to denaturation by SDS.

### 4.3. Ff-nano Functionalization and Use as a Detection Particle

To allow exploration of Ff-nano in biotechnological applications, we have converted the Ff-nano production system into a display system through insertion of multiple cloning site (MCS) into *gIII* of the helper phage/vector, between the signal sequence and the mature portion of the protein. This system was then used to develop a dipstick fibronectin detection assay using Ff-nano displaying the fibronectin-binding domain from *S. pyogenes* (Ff-nanoFnB). No unspecific binding to the analyte was detected using the Ff-nanoFnB detection particles, in contrast to residual non-specific signal observed using the full-length phage Rnano3FnB. The quantitative range of assay using Ff-nanoFnB detection particles was found to be between 1.25 ng/μL and 10 ng/μL, suitable for covering the range of Fn present in human serum. Given that variations of Fn concentration can be used as indicators of several diseases, such as bladder cancer, liver damage, defibrination syndrome, arterial thrombosis, preterm birth or ocular damage, and that these variations are within the detection range (Choate and Mosher, 1983;Cembrowski and Mosher, 1984;Weller et al., 1988;Honest et al., 2002;Mosher, 2006;Eissa et al., 2010), we believe that this assay using Ff-nanoFnB detection particles has potential as a diagnostic tool for detection of fibronectin.

In conclusion, this work describes a novel display system that functionalizes short Ff phage-derived nanorods. It further demonstrates one application for use in dipstick assays. Given a large range of publications describing applications where Ff virions are used, including the templates for assembly of inorganic structures, diagnostics, tissue templating, imaging and drug targeting (Bar et al., 2008;Petrenko, 2008;Lee et al., 2009;Souza et al., 2010;Chung et al., 2011;Dang et al., 2013;Bernard and Francis, 2014;Oh et al., 2014), Ff-nano as short non-viral functionalized particles will likely find many diverse applications.

## 6. Acknowledgements

We are grateful to Marjorie Russel for comments on the manuscript. Marjorie Russel and Peter Model are acknowledged for generously providing the Ff and P1 phage, plasmids pLS7 and pPMR132 and *E. coli* strains. We also thank Dr Dalaver Anjum for his expert technical assistance with the cryo-EM work (Pittsburgh University) and Doug Hopcroft for the TEM work at Manawatu Imaging and Microscopy Centre (Massey University). Funding for this project by a Marsden Fund Grant (contract number 02-MAU-210), Massey University Research Fund, Anonymous Donor, Palmerston North Medical Research Foundation, Massey University Institute of Fundamental Sciences Postgraduate Research Fund and the New Zealand Foundation for Research and Technology contract C03X0701, is gratefully acknowledged. S.S. was supported by a Pakistani High Education Commission Doctoral Scholarship and N.J.B. by a Massey University PhD scholarship.

